# Chromosomal position of ribosomal protein genes impacts long term evolution of *Vibrio cholerae*

**DOI:** 10.1101/2022.05.06.490600

**Authors:** Leticia Larotonda, Damien Mornico, Varun Khanna, Joaquín Bernal, Jean Marc Ghigo, Marie-Eve Val, Diego Comerci, Didier Mazel, Alfonso Soler-Bistué

## Abstract

It is unclear how gene order within the chromosome influences bacterial evolution. The genomic location of genes encoding the flow of genetic information is biased towards the replication origin (*oriC*) in fast-growing bacteria. To study the role of chromosomal location on cell physiology we relocated the *S10-spec-*_α_ locus (S10), harboring half of ribosomal protein genes, to different chromosomal positions in the fast-growing pathogen *V. cholerae*. We found that growth rate, fitness and infectivity inversely correlated the distance between S10 and *oriC*. To gain insight into the evolutionary effect of ribosomal protein genomic position, we evolved strains bearing S10 at its current *oriC*-proximal location or derivatives where the locus far from it, at the chromosomal termini. All populations increased their growth rate along the experiment regardless S10 genomic location. However, the growth rate advantage of an *oriC*-proximal location persisted along experimental evolution indicating that suppressor mutations cannot compensate S10 genomic position. An increment in biofilm forming capacity was another common trait observed along the experiment. Deep sequencing of populations showed on average 1 mutation fixed each 100 generations, mainly at genes linked to flagellum biosynthesis regulation, lipopolysaccharide synthesis, chemotaxis, biofilm and quorum sensing. We selected fast-growing clones displaying a ∼10% growth rate increment. We found that they harbored inactivating mutations at, among other sites, the flagellum master regulators *flrAB*. The introduction of these mutations into naïve *V. cholerae* strains resulted in a ∼10% increase of growth rate. Our study therefore demonstrates that the location of ribosomal protein genes conditions the evolutionary trajectory of growth rate in the long term. While genomic content is highly plastic in prokaryotes, gene order is an underestimated factor that conditions cellular physiology and lineage evolution. The lack of suppression enables artificial gene relocation for genetic circuit reprogramming.

## Introduction

In all known life forms, DNA stores the genetic information in a long polymer whose length surpasses the cell dimensions by thousands of times (1). As a consequence, the genetic material is highly packed by specific proteins that confer its structure and position within the cell (2–5). There is relatively little knowledge on the physiological role of spatial organization of the chromosome. The simplest model to tackle this issue is the bacterial cell due to its relatively small genome size and chromosome number. The bacterial chromosome is tightly organized within the confined space of the cytoplasm and along the cell cycle (6–13). This is physiologically relevant since DNA simultaneously serves as template for replication, transcription, segregation and the repair of the genetic material (1,4,14). Bacterial chromosomes have a single replication origin (*oriC*) from which DNA duplication begins bi-directionally until the terminal region (*ter*) where both replication forks meet. This configuration organizes the genome into two generally equally sized replichores along the *ori-ter* axis (6,15,16). Recent studies suggest that gene order contributes to link genome structure to cell physiology (14,16–20). The genomic position of key gene groups is widely conserved along the *ori-ter* axis following the temporal pattern of gene expression (17,18,21,22). Thus, coding sequences expressed in exponential phase are physically associated to the *oriC* region while factors transcribed in stress situations or in stationary phase cluster close to the *ter* region (17,21,23–26). Consistently, it was recently shown that some genes need a specific genomic location to achieve their function (16). In most cases, the genomic position of a gene conditions its genome-wide copy-number since those located close to *oriC* are overrepresented with respect to those close to the *ter* region during DNA replication (16,27). This differential template abundance, called gene dosage effect, leads to increase the expression of genes located in the proximity of *oriC,* although this might be more complex (28,29). The *ori-ter* template abundance gradient is particularly important in fast-growing bacteria during steady-state growth when cells display generation times shorter required time to complete genome replication. In these conditions, successive replication rounds overlap, a mechanism called Multi-Fork Replication (MFR) (16,27). In fact, many essential and highly expressed genes are clustered in this chromosomal region (6,24,30–32). In parallel, some other genes require a precise location to function, independently of their dosage (33,34). Other coding sequences, particularly those highly expressed, impact the chromosomal spatial structure (10,11) suggesting an interplay between gene order and chromosome structure(35).

*Vibrio cholerae* is a human pathogen of prime importance able to colonize the intestine causing the cholera disease that kills around 100,000 persons yearly (36). This microorganism is a common inhabitant of estuarine environments often associated to plankton (37,38). It is a highly motile bacterium thanks to a single, sheathed polar flagellum. The flagellum biogenesis depends on a regulation cascade whose master regulator is the FlrA protein (39). *V. cholerae* also alternates from a mobile to a sessile lifestyle by forming structured biofilms, a key trait for both parts of *V. cholerae* life-cycle (40–42). Importantly, flagellum-mediated swimming, chemotaxis, biofilm formation, colonization, and virulence are intimately related processes connected through complex regulatory cascades (40,43). In particular, flagellum-mediated motility and biofilm formation are inversely regulated by the concentration of the second messenger molecule cyclic di-guanosine monophosphate (c-di-GMP)(43–45). This molecule is in turn modulated by proteins carrying GGDEF and/or EAL domains displaying diguanylate cyclase and phosphodiesterase activity respectively(43–46).

*V. cholerae* displays a short doubling time (∼16 minutes) and high replication-associated gene dosage effects (24). It is also a model of bacteria bearing multiple chromosomes since it harbors a main chromosome (Chr1) of 2.96 Mbp and a 1.07-Mbp secondary replicon (Chr2) (47). Replication of both chromosomes is coordinated. Chr1 first starts its replication while Chr2 fires its replication origin (*ori2*) only after 2/3 of Chr1 are duplicated and both chromosomes complete DNA synthesis synchronously (48). Nevertheless replication-dependent gene-dosage effects exists in both replicons (30,49,50) while their relative timing modulates the gene dosage difference between both replicons. In *Vibrionaceae,* transcription and translation genes map close to the *oriC* of Chr1 (*ori1*) (31,49).

We have previously studied the chromosomal position of ribosomal protein (RP) genes to uncover the link between gene order and cell physiology (24,51). For this we manipulated the genomic location of *V. cholerae* major ribosomal protein locus *s10-spc-*α (S10), a 13 Kbp array of essential genes highly conserved among prokaryotes and also in eukaryotic organelles (52,53). By relocating S10 locus along both chromosomes (49) we generated a S10 *movant* strain set (i.e., isogenic strains where the genomic position of the locus of interest is altered) whose growth rate, infectivity and fitness decreased as a function of the distance between S10 and *ori1*. Relocation to the Chr2 showed a detrimental effect accordingly. These phenotypes were caused by the reduction of S10 genome-wide copy number that were mostly consequence of macromolecular crowding alterations rather than a deficit in the capacity to synthesize proteins (49,50,54). Further increasing S10 copy-number was deleterious our neutral. Genomic position of S10 maximizes cell physiology and fitness along both parts of *V. cholerae* lifecycle. Meanwhile, S10 is close to *ori1* among all *Vibronaceae.* Since this clade diverged from enterobacteria some 500 million year ago (55), such genome positioning is likely to be the result of selection in the long run rather than trait acquired through genetic drift. In the last years, long term evolution experiment emerged as the experimental approach of choice to directly assess evolutionary trajectories of (micro)organisms (56–59).

In this work, we analyzed how the genomic position of the major ribosomal protein locus could condition *V. cholerae* evolution in the long term. For this, we experimentally evolved strains in which the S10 locus is either placed at its original position or at the terminal region of the chromosome. We showed that the growth rate reduction observed in the latter case persisted over 1000 generations of evolution providing experimental support to the biased genomic position of these genes in fast growing bacteria. Our positional genetics approach also showed that increased biofilm formation and increased growth rate at the expense of motility is a common adaptations of *V. cholerae* to in vitro environment. Our study therefore contributes to improve our understanding of the role of gene order on bacterial cell physiology and provides insights into the long-term implication of gene order on genome evolution.

## Materials and methods

### Bacterial strains and plasmids and culture conditions

All strains derive from *V. cholerae* serotype O1 biotype El Tor strain N16961 with is Str^R^(60). Bacterial growth was done in Lysogeny Broth (LB) at 37°C agitating at 200 rpm. Strains and plasmids used in this study are listed in Table S1.

### General procedures

Genomic DNA was extracted using the GeneJET Genomic DNA Purification Kit while plasmid DNA was extracted using the GeneJET Plasmid Miniprep Kit (Thermo Scientific) or ADN Puriprep B and P (InbioHighway, Argentina). PCR assays were performed using DreamTaq, Phusion High-Fidelity PCR Master Mix (Thermo Scientific) or MINT 2X (InbioHighway, Argentina). Oligonucleotides, described in Table S2 were purchased from Macrogen, Ko.

### Experimental evolution setting

Starting from individual colonies, 3 independent cultures of each strain were propagated in 250 mL (7 generations) or 500 mL (8 generations) Erlenmeyer flasks. For each passage 1 mL of the previous flask is transferred to the next step. Populations were regularly checked for contamination by plating in LB Str. Colonies with unusual morphologies were checked phenotypically and by PCR amplifying the *V. cholerae* specific gene *rctB (3398-3407*, Table S2) or a specific region of the secondary chromosome (Tgt4_1-Tgt4_4, Table S2). Every 50 or 100 generations, the populations were frozen to build the “Frozen Fossile Record” by centrifuging 10 mL of each culture at 5000 rpm and resuspended them on 1 mL of LB supplemented with 10% of dimethyl sulfoxide (Merck). Then cultures were frozen at - 80°C until later use. (For detailed description see Supplements)

### Automated growth curve measurements

ON cultures of the indicated microorganism were diluted 1/1000 in LB. Bacterial preparations were distributed by triplicate, for evolved populations, and sextuplicates, for the ancestral clones, in 96-well microplates. Growth-curve experiments were performed using a TECAN Infinite Sunrise microplate reader, with absorbance (620nm) taken at 5-minutes intervals for 5 hs on agitation at 37°C. Experiments were done at least twice independently. Slopes during exponential phase were directly obtained using a home-made Python script coupled to growthrates program (61). To filter noise due to growth variations among experiments, the obtained µ was relativized to the ancestral clones and the relativized values were multiplied by the average of the ancestral clones along all the experiments. This normalization did not affect the results since raw data provide a similar picture as depicted in Figure S3. The % GT was calculated as follows: %GT = (GT/GT^WT^-1)*100

### Biofilm formation assays

PVC 9-well microtiter plates (BD Falcon) were used to monitor biofilm formation as described previously (62). Briefly, LB was inoculated with a 1/100 dilution directly obtained from the liquid overnight cultures of each population in rich medium. After inoculation, microtiter plates were incubated at 37□°C for 24□hs, rinsed and 150□μL of a 0.1% solution of crystal violet was added to each well. The plates were incubated at room temperature for 30□min and rinsed, and biofilm formation was tested as follows: crystal violet was solubilized by addition of 150□μl of ethanol-acetone (80:20), and the OD_570_ was determined. Results were presented as the mean of four replicate wells in three independent experiments.

### Deep sequencing analysis

The quality of Illumina reads was visualized by FastQC v0.10.1 Brabraham Bioinformatics (http://www.bioinformatics.babraham.ac.uk/projects/fastqc/). All reads of 32 samples were aligned against the two chromosomes of *Vibrio cholera* (Genbank NC_002505 and NC_002506, 17-DEC-2014), using paired end mapping mode of BWA ‘mem’ v0.7.4(63) with the option ‘-M’. The reference chromosome I was modified since the reference genome sequence (AE003852) showed an inversion around ori1 flanked by two ribosomal RNA operons (*rrnB* and *rrnG-H*) (48). Output SAM files were converted to BAM files using SAMtools v0.1.19 (64). The ‘Add Read Groups’ step was made by Picard v1.131 (http://picard.sourceforge.net/). The aligned reads in BAM files were realigned with the command ‘IndelRealigner’ implemented in GATK v2.2.16 (65). ‘MarkDuplicates’ step from Picard flagged duplicates reads. We only kept uniquely mapped reads, using SAMtools (option ‘view –bq 20’). Then, mpileup files were generated using SAMtools without BAQ adjustments (option ‘mpileup –B’). SNPs and INDELs were called by the option ‘pileup2cns’ of Varscan2 v2.3.2 (66) with a minimum depth of 10 reads, a threshold of 20 for minimum quality, a threshold of 0.25 and 0.8 for minimum variant allele frequency for heterogeneous population strains and subclone isolated strains respectively. The coverage around the mutation was checked with a python script with a minimum depth of 7 reads. The annotation of identified variants was processed by SnpEff v4.1 (67). Common mutations between all sequenced strains were ignored, indicating differences with the reference sequence. All data generated and analyzed during this study is included in this published article, its supplementary information files, and publicly available repositories. The Illumina Deep Sequencing datasets are deposited at the GenBank SRA as BioProject PRJNA816505.

### Multiplex Genome Editing by Natural Transformation (MuGENT)

To introduce the mutations observed along the evolution experiment we employed MuGENT (68,69). Briefly, the wild type strain (Table S1) was grown overnight from a single colony at 37°C in LB with agitation. This ON culture was diluted 10^-3^ into fresh medium, and the strain was grown to an OD_600nm_ ∼0.5. Cells were washed by centrifugation and resuspended at the original volume in 1× Instant Ocean Sea Salts (7 g/L) (Blacksburg) and autoclaved chitin powder (Sigma Aldrich) and incubated overnight without agitation at 30°C to induce competence. In parallel, DNA fragments harboring the mutations were synthesized using PCR assembly, cloned using pEASY^®^-Blunt Zero Cloning Kit (Transbiotech, China) and verified using Sanger sequencing (Macrogen, Ko). Next we co-transformed the targeted loci containing the desired chromosomal alterations and a spectinomycin resistance cassette directed to a neutral locus at the intergenic space between VC1903 and VC1902 region (70). After ON incubation at 30°C without shaking, the sample was vortexed and plated in LB supplemented with spectinomycin. Then several colonies were screened and confirmed by MASC-PCR (69).

### Motility Assay

The effect of mutations in *flrA* and *flrB* on the motility of *V.cholerae* was tested on LB soft agar plates (0,3% bacteriological agar). Next, 2 μL of bacterial suspension (OD_600_ = 1) was inoculated into each plate by triplicate to test bacterial swimming. The plates were incubated for 12 hs at 37 °C, then the diameter of the motility halos was measured using ImageJ (71). The results were reported in mm of bacterial extension on the media.

## Results

### *Vibrio cholerae* evolution experiments

We conducted experimental evolution of single clones of the parental or different *V. cholerae* movants for 1,000 generations. For this, we started 3 populations from isolated colonies of each strain. Due to the fast growth of *V. cholerae*, we passaged them twice a day in Lysogeny Broth (LB) supplemented with streptomycin (Str) to avoid contamination (See Suppl. Material). On one hand, we employed the parental strainS10 was not transposed (49), as reference (Populations P1, P2 and P3, Figure 1A, Table S1). We also conducted the experimental evolution of S10Tnp-35, a movant where the S10 locus was slightly relocated (P4 to P6, Figure 1A, Table S1). This movant displays no fitness nor growth rate defect (Figure 1B), it is isogenic with respect of to the rest of the movants and shows only 8 genes that are differentially expressed with respect to the parental strain (54). These populations were used as a control that any possible genetic change associated to S10 relocation *per se* could impact the evolution of *V. cholerae*. In parallel, we evolved S10Tnp-1120 (P7 to P9) and S10TnpC2+479 (P10 to P12) (Figure 1A, Table S1) where S10 was relocated to the *ter* region of Chr1 and Chr2 respectively. These are the most phenotypically affected movants displaying a ∼15% loss of maximal growth rate and lower fitness than the parental strain (Figure 1B) (50). The populations were frozen every 50 or 100 generations (Supplementary Text). We also isolated single clones at 250 and 1000 generations for later analysis.

**Figure 1:**
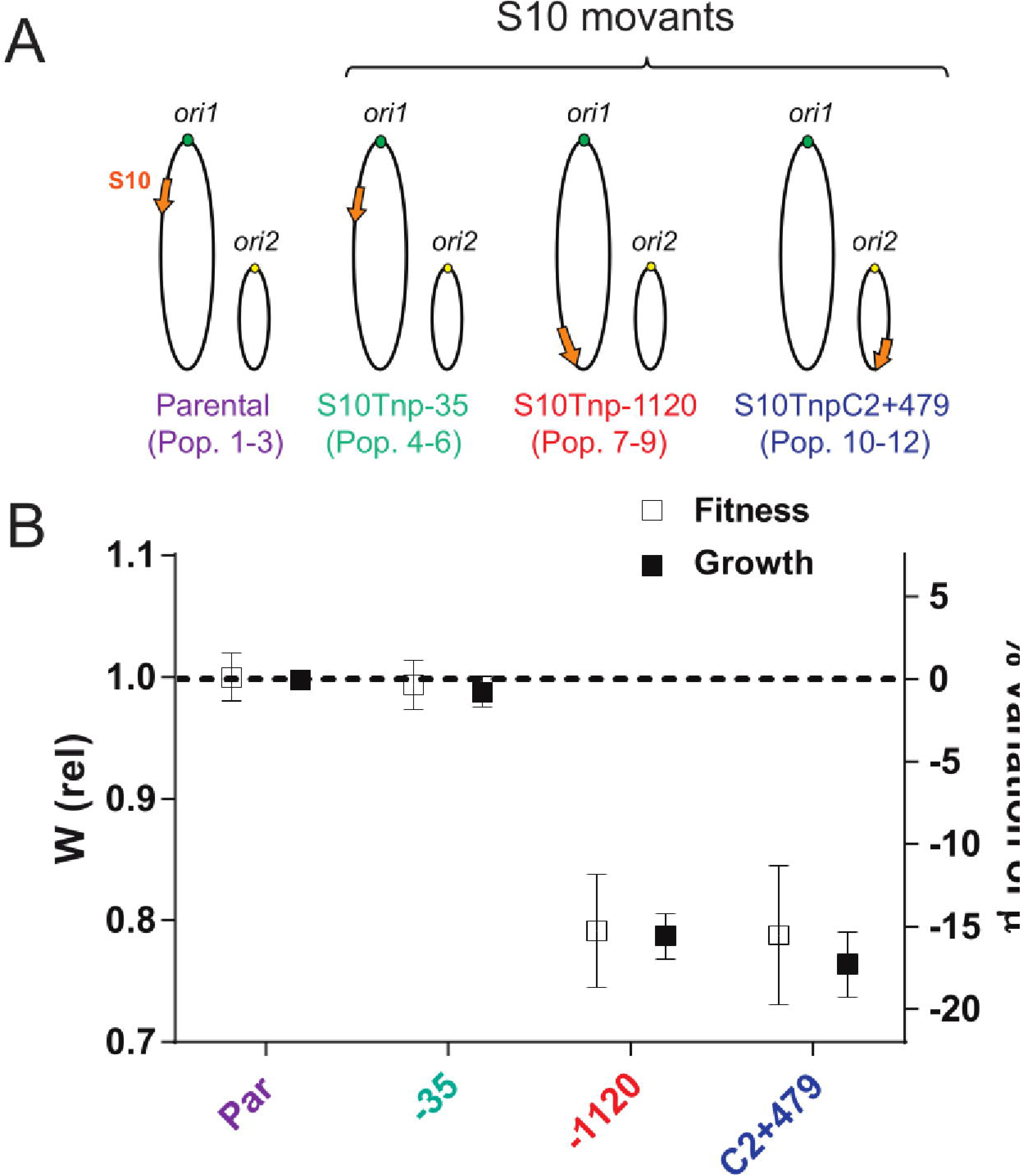
Genotype and associated phenotype of the evolved strains at the start of the experiment. **A)** Genome structure of the parental, the movants strains subjected to long term evolution for 1000 generations. Chromosome I and II are depicted as ovals-The orange arrow represents S10 at its genomic position with respect to *ori1* (green dot) or *ori2* (yellow dot). Colors are strain specific and we indicate the population numbering assigned. **B)** The variation of maximum growth rate (49)(black squares, right axis) and the relative fitness (50) (white squares, left axis) with respect to the parental strain is plotted for each strain.

### Evolved *Vibrio cholerae* populations acquire increased biofilm formation capacity

First, we noticed the emergence of strong biofilms on the Erlenmeyer flasks as early as generation 81 (G81). By generation G100, most of the populations displayed strong biofilms in the air-liquid interphase and/or dense cell aggregates in the medium (Figure S1A). By G250 biofilms were present in all populations. Accompanying this phenomenon, colonies of abnormal morphology appeared, particularly, rugose colonies (Figure S1B). The subculture of isolated smooth (S) and rugose (R) colonies showed that only the latter formed strong biofilms in the flasks (Figure S1C). To quantitatively address biofilm-forming capacity of the populations along the experimental evolution, we cultured them statically in 96-wells polystyrene plates, washed and then stained the wells with crystal violet at G0, G100, G250, G500, G700 and G1000 (Figure 2). As positive and negative controls we analyzed R1, a rugose clone isolated from P1 at G1000 (see above) and the parental strain respectively. At the beginning of the experiment (G0), the levels of crystal violet stain are minimal. Then biofilm-formation capacity increased rapidly in all populations. In average, biofilm formation capacity reached a maximum at ∼G250 (Figure 2) although some specific populations displayed an earlier peak near ∼G100 (Figure S2). After G250, the biofilm formation capacity decreased in all evolved populations without going back to the initial G0 baseline levels. The emergence of rugose colonies paralleled evolution of biofilm forming capacity (Figure S1D). All populations displayed a similar trend suggesting that it is selected along the experiment independently of S10 genomic location (Figure 2 and Figure S2).

**Figure 2:**
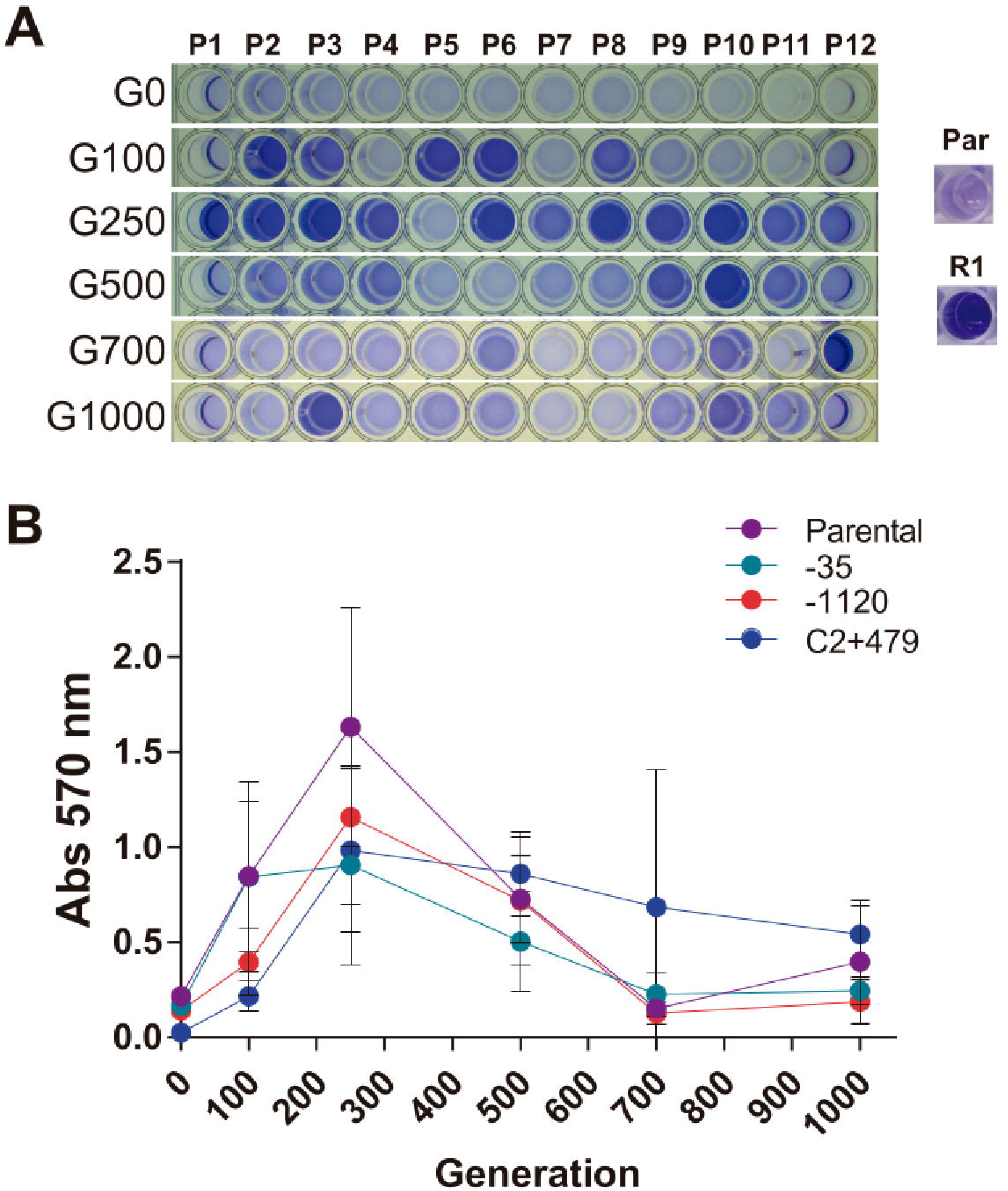
Evolution of biofilm formation capacity along the evolution experiment. Biofilm formation phenotype of twelve evolved populations from the parental strains and 3 movants was evaluated on polystyrene microtiter plates as described in methods section. **A)** Representative wells where cells were stained with crystal violet are shown for P1-P12 at different time points. **B)** The crystal violet stain was quantified for all populations along generations. The parental (Violet, P1-P3) and S10Tnp-35 (Cyan, P4-P6), S10Tnp-1120 (Red, P7-P9) and S10TnpC2+479 (Blue, P10-P12) movants populations values were averaged for simplicity (Figure S2 shows the disaggregated data). Data averaged the means from three independent experiments done in quadruplicates.

### Evolved populations display growth rate increase

Previous studies showed that bacterial populations increase their fitness along time compared to the ancestral clone (58,72). Direct fitness determination from competition experiments results is difficult to achieve in populations that form biofilms dynamically across generations. Therefore, we employed maximum growth obtained rate (μ) as a proxy for fitness estimation. In previous works, both parameters changed similarly as a function of S10 genomic location (50). Therefore, the evolved populations were subjected to automated growth curves. For this, at each time point, the twelve populations were diluted 1 in 1000 and cultured in LB at 37°C measuring OD_620nm_ by triplicate at least twice. In each experiment, the 4 ancestral strains were used as control. The value of µ was calculated from the growth curves and relativized to the parental strains to remove variations between experiments. Then we plotted the maximum growth rate as a function of elapsed generations (Figure 3A, Figure S3A). During the first 150 generations, µ remained stable among populations. As expected (49,50), populations derived strains in which S10 is close to *ori1* (P1 to P6) grew ∼15% faster than those originated from the most affected movants (P7 to P12). After this period, between G150 and G250, populations increase their µ. All populations increased their growth rate along the experimental evolution but this rise occurred in a stepwise manner, the most notorious occurring at G250. After G500, the alterations in growth rate become noisier since populations originating from the same strain tend to diverge (Figure S3A). Despite the general µ increment along generations, the differences between populations evolving from faster strains (P1-P6) and slower ones (P7-P12) persisted along the experiment (Figure 3B). Also, the evolutionary trajectory of this trait was similar for all of them (Figure S3B). Importantly, we did not notice suppressor mutants compensating the growth rate deficit in P7-P12. Hence, mutations able to compensate the genomic position of S10 locus are very difficult to achieve, at least in this time frame. This, strongly suggests that the S10 genomic location influences growth rate along experimental evolution strongly indicating that the position of the main ribosomal protein locus determines fitness in the long run.

**Figure 3:**
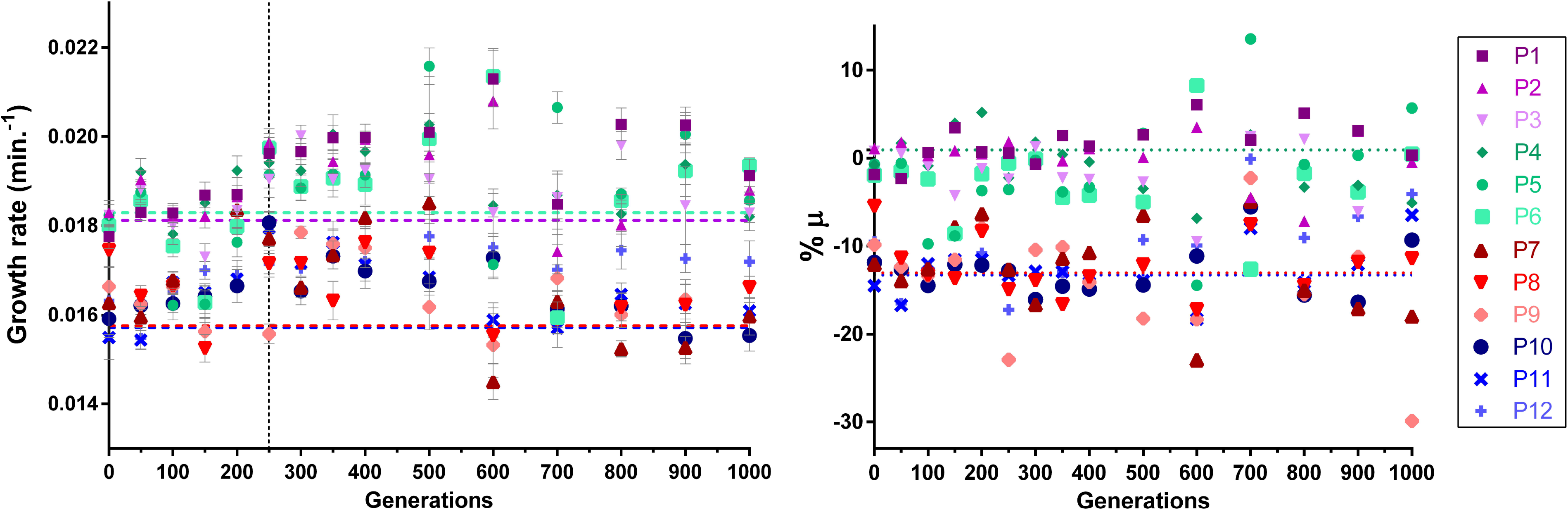
S10 genomic location impacts growth rate evolution in the long term. **A) *V. cholerae* increases its growth rate along evolution.** At every time point, the populations from the frozen “fossil record” were subjected to automated growth curve and relativized to their ancestral strains. Parentals (P1-3), S10Tnp-35 (P4-6), S10Tnp-1120 (P7-9) and S10TnpC2+479(P10-12) are shown in tones of violet, cyan, red and blue respectively. The graph shows the average at least two experiments done by triplicate of µ ± SD as a function of the generations elapsed. The average µ from the Parental (Violet) strain and S10Tnp-35 (Cyan), S10Tnp-1120 (Red) and S10TnpC2+479 (Blue) is shown as dotted lines for reference. **B) Growth rate defect of movants persists along 1000 generations of evolution.** The obtained µ for each movant population (P4-P12) was relativized to growth rate of the Parental populations (P1-P3) for each time point. Then the mean with SD of percentage of µ variation was plotted as a function of generations. The dotted lines represent the average deviation from growth of the parental for each ancestral strain (0.91% for S10Tnp-35 and −13% for S10Tnp-1120 and S10TnpC2+479).

### Positive selection drives genome dynamics in the first 250 generations of experimental evolution

To better characterize mutations occurring along the experiment, we deep sequenced each ancestral strain as reference and each population at G250 and G1000. We speculated that mutations at G250 would correlate to the earlier increase in growth rate and biofilm-formation capacity emergence.

Deep sequencing of the 12 populations at G250 revealed 128 mutations observed in more than 25 % of the reads obtained from each culture (Dataset S1). Among them, we found 45 different mutations among all the populations. The number of fixed DNA changes varied from 2 to 21 between each individual populations (Figure 4A). On average, each characterized mutation occurred in 2-3 populations strongly indicating that they were under positive selection. Most mutations (26 out of 45) corresponded to non-synonymous mutations or frameshifts of coding sequences (Figure 4B). These changes are likely to alter or interrupt their encoded functions. Meanwhile, 17 changes (42%) occurred at *a priori* non-coding regions such as intergenic spaces and pseudogenes (Figure 4B). We detected 2 (4.4%) synonymous mutations. When grouped by coding sequences (CDS) we found that at least 17 genes suffered alterations (Dataset S1). Among them, 6 genes presented more than one mutation across multiple populations: *aroG*. *ompT*, *flrA, flrB, flrC* and *tagE2* (Table 1). First, *ompT* (73) is a highly expressed gene encoding one of the main porins of *V. cholerae*. *tagE2* (VCA1043) was found altered in most of the populations. There is little evidence of its encoded function although a recent study shows it is an endopeptidase necessary for cell growth (74). The *aroG* gene is required for the synthesis of aromatic amino acids although there is little work done on *V. cholerae*. Finally, we found SNPs at *flr* genes, the main regulators of flagellum biosynthesis (39,75,76). FlrA is the master regulator of flagellum biosynthesis controlling the expression of *flrB* and *flrC* which in turn, are downstream in the flagellar transcription hierarchy (39). We reasoned that this multiplicity of independent mutational events within the genetic pathway suggests a fitness benefit from motility inactivation. As a general scenario, we found that at G250 mutations occur several times independently, suggesting a positive selection of most of them rather than genetic drift. In particular, we observed that a majority of the populations displayed mutations at the main flagellar regulators (DataSet S1, Table 1).

**Figure 4:**
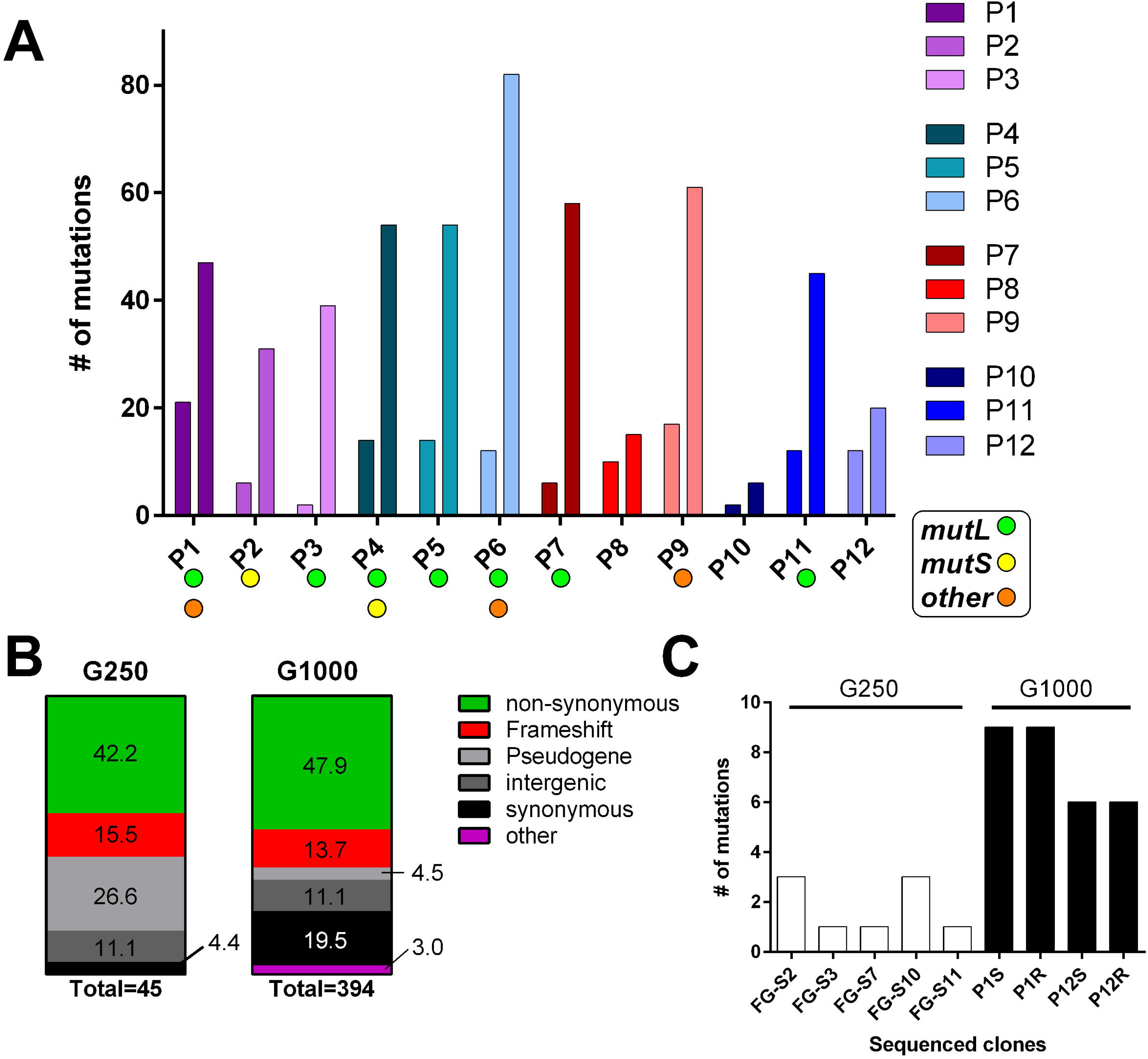
Evolutionary trajectories of the evolved populations. **A)** The graph shows the number mutations fixed to at least of 25% of the population. For each population the left and right bars indicate the mutations detected at G250 and G1000 respectively. The colors indicate the ancestral strain for the population (violet and nuances for parental; Cyan and nuances for S10Tnp-35; red and nuances for S10Tnp-1120; and blue and nuances for S10TnpC2+479). The filled circles indicate populations showing mutations at the DNA repair genes: *mutL* (green); *mutS* (yellow); others (*e.g. recBCD, recJ,* orange). **B)** Each column represent the total of the observed mutations at G250 and G1000. The colors represent the proportion of each type of mutation observed. The numbers within the color areas indicate the corresponding percentage. **C)** Total mutations fixed in the single clones sequenced. The white columns represent the fast-growing clones (FG) isolated at G250. The dark columns represent the S and R clones isolated at G1000.

**Table 1:** Recurrent mutations occurring on evolved *V. cholerae* populations after 250 generations. We included genes mutated more than once in multiple populations.

| **Gene** | **Function** | **Population** | **Type of mutation<sup>1</sup>** |
| --- | --- | --- | --- |
| *ompT*<br>(VC1854) | Main porin | 9,10 ,11 and 12 | FS, S' |
| *flrA* (VC2137) | Flagellar master regulator | 1, 6 | FS, NS |
| *flrB* (VC2136) | Flagellar regulator (Class II) | 3, 4 | FS, NS |
| *flrC* (VC2135) | Flagellar regulator (Class II) | 2, 9, 10 | FS, NS |
| *aroG*<br>(VCA1036) | Aromatic amino acid synthesis | 1, 6, 8, 9, 12 | NS |
| *tagE2*<br>(VCA1043) | Endopeptidase | 1,4,5,7,9,11,12 | NS |

<sup>1</sup>S: Synonymous mutation; NS: non-synonymous mutation; FS: Frame-shift mutation.

### Evolutionary forces change beyond G250: adaptative, non-adaptative mutations and hypermutators after 1000 generations of experimental evolution

We speculated that mutations observed at G1000 may reflect those that are fixed in the long run since after ∼G500 biofilm formation decreases and growth rate remains roughly constant. After 1000 generations of experimental evolution, we observed 512 fixed changes in at least 25% of the population at 396 sites of the genome of *V. cholerae* (Figure 4A, right bars). Contrasting to G250 scenario, few mutational events occurred in more than one population (1.3 events per populations on average). We observed a lower proportion of mutated pseudogenes (4.5% vs 26.6%) and an increase in the proportion of synonymous mutations (4.4% vs 19.4%) (Figure 4B). We also noticed mutational events at several genes involved in DNA repair such as *mutT* (VC1342), *recBCD* (VC2319-VC2320), *mutS* (VC0535), *recJ* (VC2417). Particularly, *mutL* (VC0345) was mutated in 8 out of 12 populations (1, 2, 3, 4, 5, 6, 7 and 11). Indeed, populations with intact DNA repair mechanisms (i.e. P8, P10; P12) are those showing lowest number of mutations (Figure 4A). These data suggest that in most populations at G1000, many of the identified mutations may result from an hypermutator phenotype rather than selection. All, fast-growing populations (P1 to P6) showed one or more mutations in DNA repair systems. In contrast half of the slower-growing populations (P7 to P12) displayed an intact DNA repair pathway (Figure 4A). Nevertheless, we also detected many DNA changes in the same gene or operon occurring many times independently or emerging in parallel in more than one population likely indicating positive fitness selection along the evolution experiment (Table 2). On top of *aroG, ompT*, *flrABC* and *tagE2* already acquired at G250, we found many mutations in interlinked cellular processes (38,43,44) such as rugosity, biofilm structure and regulation (*vpsA, vpsB, vpsP, vpsR, capK and rbmB*) (77–80), quorum-sensing (*luxP, luxO* and *luxU*)(81), flagellum biosynthesis (*flgA, flgI, flgK, fliG, fliO, flag, flhA*)(39) and chemotaxis (*cheY, cheR2*) (39,82). We also identified several altered functions such as extracellular macromolecular structures linked to virulence (toxin coregulated pilus (*tcp*), mannose-sensitive hemagglutinin pilus (MSHA), type 6 secretion system (T6SS)), iron incorporation and metabolism (particularly vibriobactin pathway genes and *iscR*), lipopolysaccharide biosynthesis (*rfbA, rfbB, rfbV, pgi*) and phosphate sensing and incorporation (*phoR, phoU, phoH*). Interestingly, the latter have been linked to biofilm downregulation (83). Most of them were non-synonymous or led to frameshifts likely to be detrimental for their biological function (Data Set S2). We interpret these mutations in the light of a change from a host-environmental life cycle to the relatively constant culture condition of the evolution experiment. Functions such as virulence, swimming, chemotaxis, pili production, T6SS and micronutrient scavenging are energetically costly structures under strong selection in *V. cholerae* natural environment that may become a burden during *in vitro* evolution.

**Table 2:** Mutations occurring in multiple populations of evolved *V. cholerae* populations after 1000 generations. All mutations recovered and the details are described in Dataset S2.

| Gene/ORF<sup>1</sup> | Function | Populations<sup>2</sup> | Type of mutation<sup>3</sup> |
| --- | --- | --- | --- |
| *rfbB* (VC0242) | Phospho-mannose mutase | 2, 3 | NS |
| *rfbV* (VC0259) | Lipopolysaccharide biosynthesis | 2, 5 | NS |
| *manA-1* (VC0269) | Lipopolysaccharide biosynthesis Mannose-6-phosphate isomerase | 1, 3, 4, 6, 7, 8, 9, 11 | FS |
| intergenic (VC0341(orn),tR NAGly) | Unknown | 1, 6 | - |
| *mutL* (VC0345) | Mismatch repair | 1, 3, 4, 5, 6, 7, 11 | NS, FS |
| *mshH*(VC0398) | MSHA biogenesis | 1, 9 | NS |
| intergenic tRNA33-VC0449 | Unknown | 1, 5, 8 | - |
| intergenic VC0511-12 | Unknown | 4, 7, 8, 9, 12 | - |
| *mutS* (VC0535) | Missmatch repair | 2, 4 | FS |
| VC0590 | Permease | 7, 11 | NS |
| VC0603 | Unknown (PNPLA domain-containing protein) | 1, 5, 6, 7, 11 | FS |
| *iscR* (VC0747) | Fe-S cluster sensor status and biogenesis | 1, 3, 4, 5, 6, 9, 10, 11 | NS,S,FS |
| *tcpC* (VC0831) | Toxin coregulated pilus | 1, 3, 6, 7, 9 | FS |
| *gcvA* (VC0896) | Transcriptional activation of the glycine cleavage system. | 4, 9 | NS |
| *vpsA* (VC0917) | Biofilm formation | 2, 6 | NS |
| *luxO* (VC1021) | Quorum sensing regulation, Biofilm formation | 1, 2, 3, 5, 6, 10 | NS |
| VC1724 | Hypothetical protein | 5, 6 | FS |
| VC1761 | Hypotetical protein (VPI-2) | 4, 9 | NS |
| *ompT* (VC1854) | Main porin | 11, 12 | NS,S,FS |
| *flhA*(VC2069) | Flagellar biosynthesis protein | 1, 6 | FS, NS |
| VC2105 | Putative Thioesterase | 1, 4, 5, 7, 9, 12 | NS |
| *flrA* | Flagellar master regulator | 1, 2, 3, 4, 5, 6, 7, 9, 10, 11, 12 | FS, NS |
| Intergenic (VC2286-VC2887) | Unknown | 2, 3, 9 | - |
| Intergenic (VC2387-2388) | Unknown | 1, 6 | - |
| *rimO* (VC2620) | Ribosome biogenesis | 1, 6, 7 | FS, NS |
| VC2684 | Bifunctional aspartokinase/homoserine dehydrogenase | 7, 10 | NS |
| VC2707 | Unknown | 1, 9, 12 | NS |
| VCA0044<br>(Pseudogene) | Diguanylate cyclase | 8, 12 | NS |
| VCA0088<br>(Pseudogene) | Proton/glutamate symporter | 6, 7, 12 | NS |
| VCA0168<br>(pseudogene) | BatD family protein | 1,2, 4 6, 7, 8 ,9 ,11, 12 | - |
| VCA0354<br>(Pseudogene) | DedA family protein | 6, 9, 12 | - |
| Intergenic<br>(VCA0591-VCA0592) | Unknown | 6, 9 | - |
| *luxP* (VCA0737) | Quorum Sensing | 3, 7, 9 | NS |
| VCA1031<br>(Pseudogene) | Chemotaxis Like protein<br>(10.1016/j.femsle.2004.08.03<br>9) | 2, 4, 5, 6, 7, 8, 9, 11, 12 | - |
| *tagE2*<br>(VCA1043) | Endopeptidase | 1, 4, 5, 7, 8, 9, 12 | NS,FS |
| intergenic<br>(VCA1052-3) | Unknown | 1, 3, 4, 5, 7 | - |
| *aroG* (VCA1036) | Aromatic amino acid synthesis | 5, 7 | NS |
| VCA1080 | HlyD family secretion<br>protein | 1, 2, 6, 8 | NS |

<sup>1</sup>We included intergenic regions specifying the flanking genes <sup>2</sup>S: Synonymous mutation; NS: non-synonymous mutation; FS: Frame-shift mutation.

### Mutations linked to rugose phenotype

Along the experimental evolution we observed the emergence of biofilms and aggregates linked to a rugose phenotype described earlier (Figures 2, S1 and S2). To uncover genomic changes associated to biofilm emergence we fully sequenced the genome of a rugose and a smooth clone from a slow growing (P1) and a fast-growing population (P12). After 1000 generations, clones originated from P1 and P12 fixed 9 and 6 mutations respectively (Figure 4C) independently of their smooth (S) or rugose (R) phenotype (1 mutation fixed every 133±30 generations). Although some DNA changes were common among R and S colonies (Data Set S3), we found some differential mutations potentially involved in biofilm and rugose phenotype emergence (Table 3). In particular, mutations in *vpsR, vpsB, cdgB* and VCA1031 were previously linked to rugose phenotype by Yildiz and co-workers (46,61,84). The *vpsR* is a positive regulator of the *vps cluster* that suffered a non-synonymous mutation (D40Y) in the S colony derived from P1. The *vpsB* gene codes for an enzyme participating in the synthesis of nucleotide sugar precursor necessary for polysaccharide biosynthesis. We detected a premature stop codon at the S colony collected from P12 (P12S). The P12R displayed a non-synonymous mutation (D101G) at the *cdgB,* a GGDEF protein upregulated in rugose clones (61,85). It also showed a non-synonymous mutation in the *iscR* gene which codes for a transcription factor regulating genes related to iron starvation, oxidative stress and oxygen limitation in *Escherichia coli* and linked to virulence in other bacteria (86). Meanwhile, the P1S colony displayed a mutation at VCA1031, encoding a putative methyl-accepting chemotaxis proteins previously reported to be overexpressed in rugose strains having a genetic *ΔcdgC* background(46). In parallel, P1R showed an intergenic mutation also found at G250, G1000 and on fast-growing clones (see above). Finally, P12S displayed a synonymous mutation in the *rpoA* gene, a main component of the RNA polymerase. We found two other differential mutations unlikely to be causal rugose phenotype but that might have been co-selected. In summary, the S and R clones showed alterations in genes previously linked to biofilm generation and c-di-GMP regulation. We detected, however, other changes in R clones such as in *iscR* and at the intergenic region between VC0448-VC0449.

**Table 3:** Mutations linked to rugose phenotype on *V. cholerae* populations evolved for 1000 generations. Dataset 3 includes the full list of mutations and their details.

| Gene | Function | Population | Type of mutation<sup>1</sup> |
| --- | --- | --- | --- |
| *vpsR* (VC0665) | Positive Biofilm regulator | 1S | NS |
| *vpsB* (VC0918) | Biofilm | 12S | NS |
| *cdgB* (VC1029) | GGDEF protein | 12R | NS |
| VCA1031<br>(Pseudogene) | Chemotaxis Like protein | 1S | NS |
| *iscR* (VC0747) | Fe-S cluster sensor status and biogenesis | 12R | NS |
| *rpoA* (VC2571) | $\alpha$ subunit of RNA polymerase | 12S | S |
| Intergenic<br>(VC0341(*orn*),<br>tRNA-Gly | Unknown | 1R | - |

<sup>1</sup> S: Synonymous mutation; NS: non-synonymous mutation; FS: Frame-shift mutation.

### Increase growth rate is often associated to motility loss

To uncover mutations associated to fast growth, we isolated 4 smooth colonies and a rugose clone (S1 to S4 and R respectively) from P1 to P12 at G250 the moment along experimental evolution where we noticed a sudden growth rate increase in all populations. Next, we performed automated growth curves of these clones using as reference the parental strain Par-1120 (Table S1). We calculated µ for each strain (Figure 5A) and we selected the fastest growing clones from P1 to P6 and from P7 to P12. We called these clones <u>F</u>ast <u>G</u>rowing clones followed by population of origin and its smooth or rugose status. Thus, we isolated the clones FG-2-S2, FG-3-S2, FG-7-S1, FG-10-S1 and FG-11-S1. Interestingly, the 3 former clones grew 7 to 10% faster than the parental strain. Meanwhile, FG-G250-7-S1, FG-G250-10-S1 and FG-G250-11-S1 improved their growth rate compared to their ancestors, they still displayed a µ reductions of ∼7% with respect to the parental strain (Figure 5B). To identify the mutations responsible of µ increase, we fully sequenced FG-G250-2-S2, FG-G250-3-S2, FG-G250-7-S1, FG-G250-10-S1 and FG-G250-11-S1 genomes and compared them to the ancestral strains. Overall, strains displayed 1 to 3 changes in the DNA sequence (1 mutation fixed every ∼140 generations) (Figure 4C). In the 5 strains we found 8 different mutations (Table 4). All the isolated strains harbored alterations either at *flrA* (4 out of 5) or at *flrB,* (1 strain) the main flagellum regulators which were previously observed at the population level at G250 (Table 1). Three other mutations accompanied the latter: a non-silent mutation at a magnesium transporter (*mgtE*), a point mutation at an intergenic base between genes VC0448 and VC0449 and a frameshift mutation at *ompT*. Except for the latter, we found the exact same mutations at G250 and most of them at G1000 (Dataset S4).

**Figure 5:**
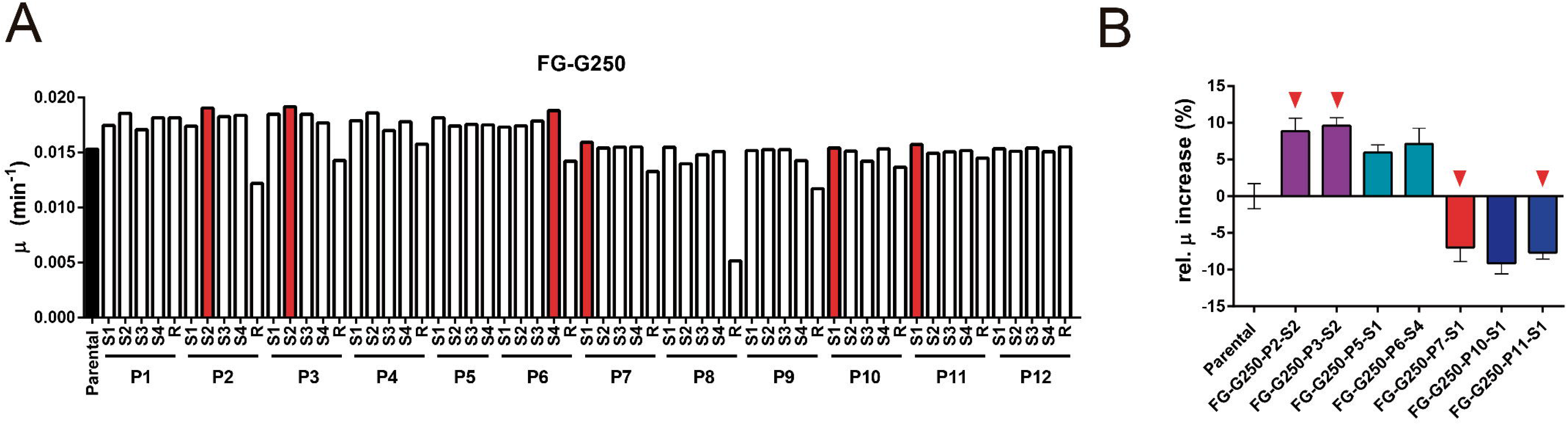
Specific mutations associated with fast growth rate. **A)** Screening fast-growing clones: 5 isolated clones per population (four S and one R) were tested per triplicate. The graph shows the average µ of each clones compared to Parental-strain (black). Clones indicated in red were selected for further study. **B)** The clones selected in A) were subjected to growth curves and the % variation of µ with respect to the parental strain is shown. The indicated clones were subjected to Deep sequencing. **C)** The mutations identified in clones from B were introduced in a parental naïve genetic background.

**Table 4:**
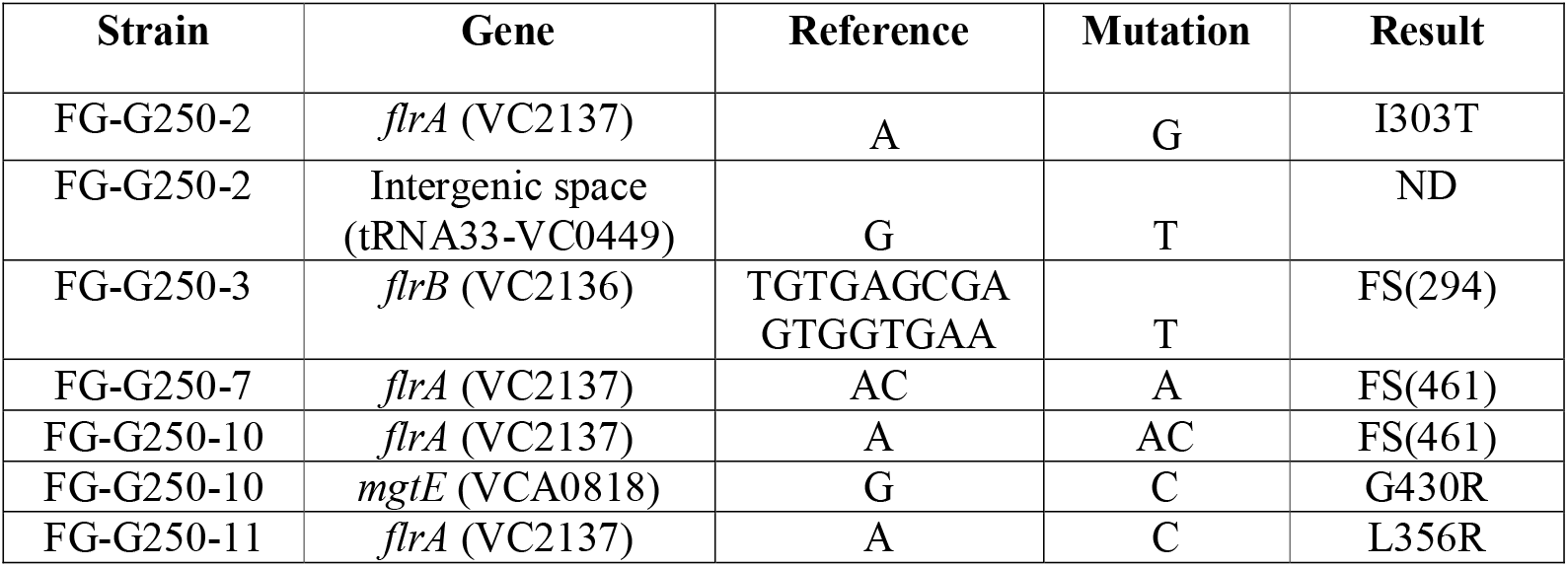
Mutations observed in fast-growing clones isolated from G250. Details are described in Dataset S4.

To demonstrate the role of these mutations on growth rate enhancement we performed Multiplex Genome Editing by Natural Transformation (MuGENT) (68,69) on naïve *V. cholerae* N16961 to introduce point mutations, small indels or short insertions. *flrA* is higher in the hierarchy of the flagellum regulation (39) and it is more prevalent on FG strains. FlrA is a 488 aminoacid-long protein bearing a REC domain, a central ATPase Associated with diverse cellular Activities (AAA+) and a C-terminal Helix-Turn-Helix domain responsible for DNA binding (87). The detected *flrA* frameshifts mutations on populations or on FG-G250 clones occurred exclusively at the end of this latter domain, on positions 418-488 of the protein. The non-synonymous mutations occurred at the AAA+ domain (positions 122 to 417 of FlrA). Using a *V. cholerae* wild type background, we generated a frameshift at the same in *flrA* position (Fig S4) obtaining several independent mutant clones that we tested on automated growth curves (Figure 6A). This mutation increased growth rate with respect to wild type strain. Similarly, the introduction of a frameshift in *flrB* resulted in a growth rate improvement. Also, the introduction of a non-synonymous mutation at *mgtE* significantly improved growth rate (Fig 6A). To quantify the effect of individual mutations we determined the percentage of generation time variation (%GT). Mutants bearing the *flrA* frameshift displayed the strongest reduction with a 13.41±4.7 % reduction of their generation time (95% CI [14.9-11.9], p<0.0001). Meanwhile, *flrB* and *mgtE* mutants displayed a 6% of generation time reduction (95% CI of [6.9-5.4] and [7.5-4.6] respectively, p<0.0001). These results coincide with the observed increase in growth rate of FG clones. Simultaneously, frameshifts in *flrA* and *flrB* caused a reduction of motility as shown by swimming assays (Fig. 6B) demonstrating that growth rate increase is associated to the loss of *flrA* and *flrB* function.

**Figure 6:**
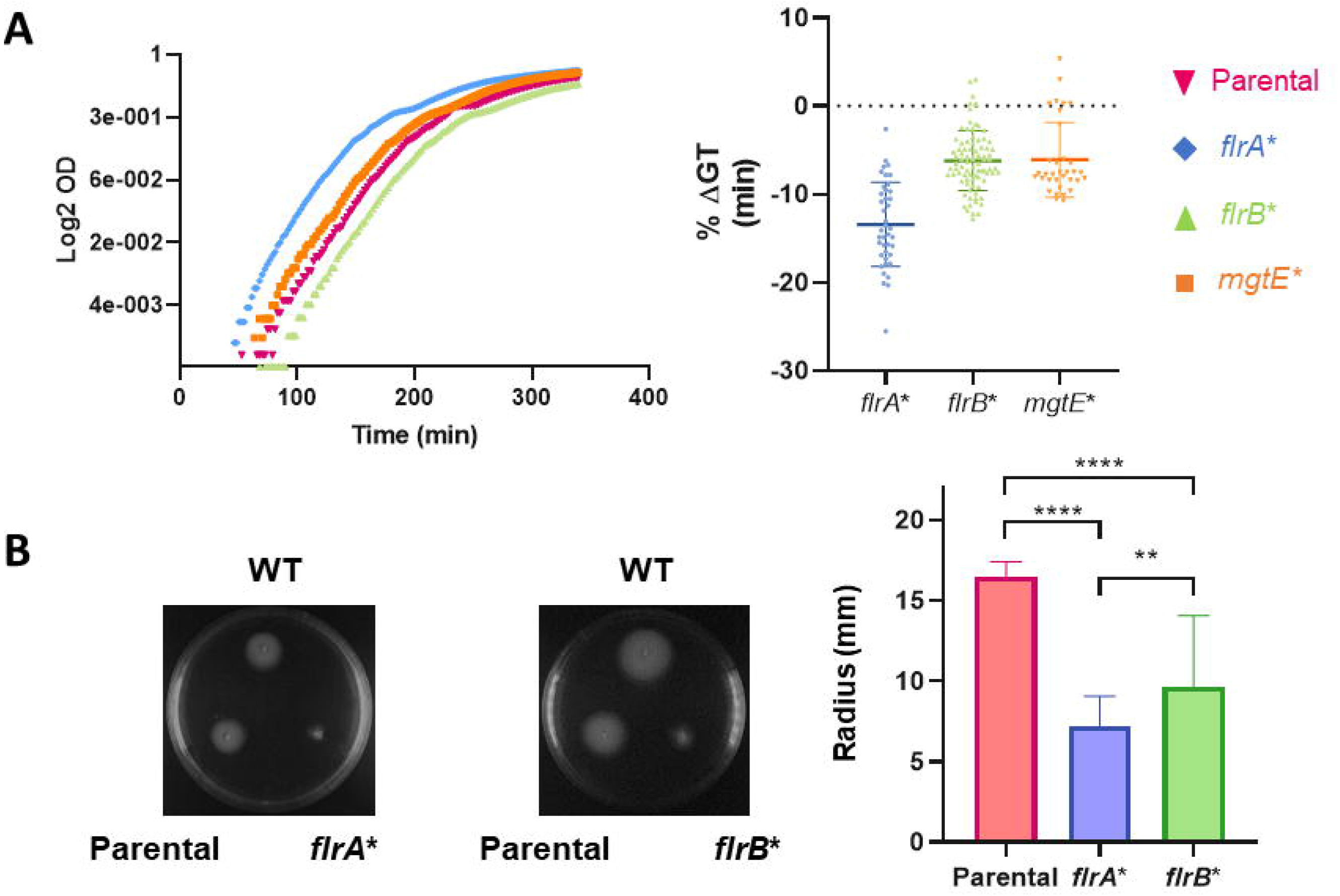
Fast-growth associated mutations increase growth rate in a *V. cholerae* naïve context. **A)** Mutations associated with fast growth (Table 4) were introduced wild type *V. cholerae* using MuGENT. The growth rate of these mutants was determined in automated growth curves performed by triplicate (n=11). Left panel shows a representative growth curve. Right panel represents the median ± CI of the % generation time (GT) variation of the indicated mutants with respect to the wild type (WT). Statistical significance was analyzed by one-way ANOVA. Then Dunnet’s test was done for multiple comparisons. **B)** Mutations in *flrA* and *flrB* increasing growth rate reduce *V. cholerae* swimming capacity. Left panel show a representative swimming assay of the WT, the parental strain bearing the selection cassette (Parental) and the clone bearing the indicated mutation (*flrA*\* and *flrB\**). The right panel shows the analyzed data for all the assays performed per triplicate (n=9). Statistical significance was analyzed by one-way ANOVA. Then Tukeýs test was done to compare the mean values. ** p<0.01 and **** p<0.0001.

In summary, these results demonstrate that the mutations observed in the FG strains increased growth rate in *V. cholerae*. They also suggest some of the mutations detected at G250 and maybe at G1000 also contributed to the growth optimization of this microorganism.

## Discussion

Increasing evidence suggests that gene order within the bacterial chromosome contributes to cellular homeostasis by coordinating key entangled processes such as chromosome structuration, cell cycle, replication and the expression of genetic information. While genome reshuffling experiments have not been yet performed in bacterial systems (88) approaches such as comparative genomics (17,18,21,23,31,89), systems biology (20), large DNA inversions (90–92) and relocation of individual gene sets (16,19,93–95) have provided support to this notion. Genes encoding for the genetic information flow are interesting models to test the role of genomic position on cellular physiology. The chromosomal position of rRNA, some tRNAs, RP and RNA polymerase genes is biased towards the *oriC*, particularly in fast-growing bacteria (17,22,24,51). Relocation of S10, the main RP locus harboring half of its encoding genes, far from *oriC* constrains bacterial growth (49). Competition experiments of S10 movants against wild type strains suggested that RP genomic position implied a strong fitness cost that must had evolutionary consequences (50). In this work, we took this “positional genetics” approach (in which genes of interest are systematically relocated within the genome) a step further by observing how movants bearing S10 at different genomic locations evolve *in vitro* for 1000 generations in fast growing conditions, those in which phenotypic effects were stronger (Figure 1).

### Comparison with previous experimental evolution approaches

Pioneer work by Richard Lenski and co-workers on long term evolution experiment was done for many more generations than ours (i.e. more than 75,000 (and counting!) for *E. coli*) however the strongest fitness improvement occurred during the first 2,000 generations (58,72). Therefore, although we performed a significantly shorter experiment, we believe that we observed a representative part of the evolutionary trajectory. Meanwhile we developed a different approach: we used rich media and a model organism that grows almost twice as fast as *E. coli*. On the other hand, our system used 1 log larger bottlenecks that previous studies improving the chance of mutant selection and lowering the effects of genetic drift (i.e. we transferred 10^9^ instead of 10^8^ CFU). Also we evolved populations with same genotype but bearing the key locus S10 at different genomic locations.

### Biofilm dynamics along *V. cholerae* evolution

Along this evolution experiment we found the emergence of abundant biofilms and aggregates in every population. As in prior studies (61,79,80,84), this phenotype was linked to the appearance of colonies of rugose morphology. Usually there are different types of rugose phenotypes in *V. cholerae* (77), but exploring their emergence along the experimental evolution was beyond the objectives of our study. Other experimental evolution approaches also led to biofilms and/or aggregates. In *Acinetobacter baylyii* ADP1 (96) it was the consequence of IS action that led to evolve aggregating clones. The studies on *Burkholderia cenocepacia* (97) and *Bacillus subtilis* (98) actively selected for biofilm emergence with a wide variety of clones with clear fitness advantages such as labor division and heterogenicity emergence within the structure. In our system, these traits emerged rapidly in all populations. Then, biofilm formation was reduced along the generations (Figure 2). Such dynamic behavior may reflect its selection by association to other traits such as flagellum loss, followed by decline (see above), the differentiation of more specialized sub-populations that are more efficient to specific microenvironments (such as air-liquid interphase), the frequent but stochastic emergence of this trait or the beneficial nature of this characteristic early along *in vitro* evolution. These possibilities are not exclusive. Meanwhile, we also characterized some mutations that could explain the emergence of rugose clones associated to biofilms. A limitation of our study is that we sequenced rugose clones at G1000 instead of G250 when most of biofilms are observed. At this timepoint, many other unrelated mutations could be co-selected with the DNA changes causing rugose phenotypes. This is reinforced by the fact that, similarly to previous work, we found mutations in several DNA repair systems. Whereas, understanding biofilm development was not the primary goal of our work, we described its emergence along the experimental evolution and we observed some mutations co-occurring with this phenotype that are in agreement with previous seminal papers by Yilditz and co-workers (38,78–80,84,85).

### Growth rate evolution along generations

Another clear phenotype that arose along the experiment was growth improvement of all *V. cholerae* populations (Figure 3). This trait was not initially noticed in the long term evolution experiment performed in *E. coli* by Lenski and co-workers. Meanwhile, fitness and cell size increment was observed since the initial steps of the experiment (72,99,100). Only recently, an increase in growth rate was also measured (101) which is in concordance with previous studies that show a close correlation between growth rate, cell size and fitness (102). In *V. cholerae* experimental evolution, the reduction of populations doubling times was clearer than in *E. coli* since growth rate could be detected using automated high-throughput growth measurements. This behavior could be the result of employing a richer broth, the evolution strategy (wider bottlenecks), and/or the faster growth nature of *V. cholerae*. Meanwhile, growth rate is an acceptable proxy for fitness particularly in our system as we observed in previous work (50,103,104). Therefore we propose that the increment in growth rate observed along *V. cholerae* experimental evolution reflects, at least in part, the expected fitness improvement. The stabilization of growth after G500 can be the result of high divergence between populations or the consequence of terminating the experiment relatively early, since further generations accumulation probably should lead to new increases in growth rate in a stepwise manner.

### Two phases in the evolution dynamics of *V. cholerae* genome

Deep sequencing of the ancestral clones and the populations at G250 and at the end of the experiment suggested different selection forces at each time point. At G250 each mutation occurred in parallel in ∼3 populations strongly suggesting a positive selection. Since these populations displayed a sudden gain in growth rate and biofilm forming capacity, we could associate these mutations (Table 1 and DataSet1) to the observed phenotypes (Figs. 2 and 3). This scenario changes at G1000 since we find that most mutations are represented in single populations suggesting that most of them arise by chance not being under strong selection. This is reinforced by the fact that many populations show inactivation of DNA repair systems. As expected, populations displaying mutations in repair systems resulted on average on more DNA changes after 1000 generations of evolution (Figure 4a). Interestingly, populations evolved from faster strains (P1-6) displayed more alteration in repair systems and concurrently more mutations at G1000. We speculate that due to their faster growth rate P1-P6 spent more time at stationary phase along the experimental evolution. These repeated stress situations might have increased the chances of mutator phenotype emergence.

Some DNA changes arose in more than one flask in particular inactivating genes encoding functions that might become unnecessarily costly during in vitro continuous culture (i.e. virulence, motility and nutrient scavenging). Interestingly, those acquired at G250 are not lost suggesting their continuous selection along the experiment. Moreover, in the case of flagellum biosynthetic pathway we found mutations in transcriptional regulators of lower hierarchy such as *flhA* or components of the machinery such as *flgA*, *flgI* or *flgK*. We also noticed several mutations at quorum-sensing and chemotaxis-related genes at G1000 (see above). The former seems to be a common feature of laboratory domesticated *V. cholerae* strains as previously described by Blockesh Laboratory (105).

At both timepoints, we detected several mutations occurring at *a priori* non-coding regions of the chromosome such as intergenic regions and pseudogenes. We cannot discard a biological effects through interfering promoter or regulatory regions, potential uncharacterized RNA genes or pseudogenes expressing truncated but functional proteins. This might be the case of those occurring in several populations independently such as the point mutations occurring at VCA0168, a pseudogene located in the secondary chromosome, mutated in 9 out of 12 populations (Dataset S1-S3). Interestingly, VCA0168 is actively transcribed from an upstream transcription start site. It forms an operon with VCA0169 and VCA0170 that is conserved among other *V. cholerae* strains.

### Mutations further increasing *V. cholerae* growth

Importantly, we were able to distinguish mutations that improved bacterial growth from those increasing biofilm-forming capacity. Notably, the master regulator for flagellum biosynthesis, *flrA,* and *flrB* downstream in the same regulation cascade, displayed mutations at G250 and on the FG smooth clones. These genes were intact in rugose clones isolated at G1000 indicating that we could discriminate between DNA changes improving growth rate from those involved on biofilm formation (Fig. 5 and Table 4). In the case of *flrA*, we detected 3 non synonymous mutations and two frameshifts occurring only in the last 20-30 amino acids of this 488 codon ORF at the HTH domain. Mutations at *flrA* emerged in 4 and 11 out of 12 populations at G250 and G1000 respectively. In most cases, these DNA alterations led to the described frameshifts (Dataset S1, S2 and S4). The introduction of a similar frameshifts into *flrA* and *flrB* in a wild type background increased *V. cholerae* growth demonstrating its effect on growth and strongly suggesting that the rest of the detected mutations contributed to the enhanced growth of the populations (Fig 6A). Both mutations also caused a reduction in motility (Fig 6B) indicating that these mutations alter the DNA binding capacity of FlrA and FlrB. Recent studies from Sourjik and colleagues provide an ecological and physiological interpretation of the genes that mutated along our experimental evolution (106–108). Their work pointed out a clear trade-off between motility and growth rate. In changing environmental conditions, cells deal several simultaneous energetically costly processes such as growth, motility chemotaxis and metabolism. Energy allocation to flagellar genes production, rotation of flagellum and, to a lesser extent chemotaxis, significantly reduces growth in *E. coli* (107,108). It is advantageous to dedicate energy to motility and chemotaxis in poor and heterogenous environments. The FG-G250 clones displayed mutations on the main regulators of flagella (*flrAB*) which permit the allocation of more resources for growth outcompeting wild type strains. Consistently, at G1000 we also noticed many mutations related to other swimming and chemotaxis genes (e.g. *fliO, flrC, cheY, cheR2, flgA*). In summary, many of the mutations detected along our experiment alter the motility/growth interplay in all the studied populations. However many other mutations that contribute to improve the fitness remain to be better characterized. For instance, mutations on *mgtE*, which is not clearly related to motility also improved growth rate. These opens the possibility of finding other mutations on genes not related to motility capable of improving the growth of an already very fast growth bacterium.

### Conclusions

In this study, we showed that the genomic location of ribosomal proteins conditions cell physiology in the long term. Growth impairment caused by ectopic location of the S10 locus was not suppressed by mutations elsewhere in the genome. After 1000 generations of experimental evolution, although all populations shortened their generation time, those harboring S10 locus close to *ori1* remained the fastest. This evidences the great potential of positional genetics approach (49) as a Synthetic Biology tool to condition cell physiology in a permanent avoiding the escape of suppressor variants. The latter conditions the design of stable genetic circuits. For instance, recent work by Izard *et al*. (109) created a growth switch in *E. coli* by controlling RNA polymerase expression from an inducible promoter than required 3 copies of the main repressor to avoid regulation escape. Despite the powerful nature of this synthetic circuit some escape could still be detected. Positional genetics offer a new way to cope with this problem to generate robust genetic circuits immune to suppressor mutations.

By applying the positional genetics approach on ribosomal protein genes, we contributed to understand the role of gene order on bacterial cell physiology. The present work incorporates the evolutionary aspect to studies on positional genetics expanding our knowledge on the long-term implication of gene order on genome evolution. The role of the genomic position of other key genes remain to be explored such as ribosomal RNAs(110), tRNA^UBI^(22,24), RNA polymerase(51), ATP synthase(17), among others. This is central to understand how genome’s primary structure impacts cell physiology, in particular growth rate and genome evolution. Such knowledge will enable bacterial growth repurposing while allowing to understand the behavior of more complex biological systems thus promising a deep impact in genome design, bioengineering and biotechnology.

## Acknowledgements

We are grateful to Eduardo P. Rocha, José Antonio Escudero, Alexandra Nivina, Céline Loot, Soledad Guidolín, Lucas Saposnik and Inés Marchesini for useful discussions. We thank the technical assistance of Laurence Ma and Christiane Bouchier from the Institut Pasteur Genomics Platform for DNA sequencing. We thank Pierre Faure and Julie Lambert for their technical help.

