## Supplementary material for "Chromosomal position of ribosomal protein genes impacts long term evolution of *Vibrio cholerae*": Table S1

**Table S1.** Full list of plasmids, bacteria strains used in this study:

| **Name** | **Relevant genotype and/or phenotype** | **Reference** |
| --- | --- | --- |
| ***Escherichia coli*** | | |
| XL-Blue | *recA1 endA1 gyrA96 thi-1 hsdR17 supE44 relA1 lac* [F´ *proAB lacI^q^ Z∆M15* Tn10 (Tet^R^ )] |  |
| ***Vibrio cholerae*** | | |
| Wild type | N16961::mTn*7hapR^+^* Δ*lacZ*. Er^S^, Gn^R^ and Cm^S^. | Val et al. 2012 |
| Parental | PGB-A192::*attB*’*-lox66-dfrB1-lox71* inserted in the intergenic region between VC1508-VC1509. Er^S^, Gn^R^ and Cm^R^. | Soler-Bistué et al. 2015 |
| S10Tnp-35 | S10 relocated next to its original location in the intergenic region between VC2536-VC2537. | Soler-Bistué et al. 2015 |
| S10Tnp-1120 | S10 relocated near the *dif* region of chromosome 1 in the intergenic region VC1508-VC1509. Er^S^, Gn^R^ and Cm^R^. | Soler-Bistué et al. 2015 |
| S10TnpC2+479 | S10 relocated near the *dif* sequence of chromosome 2 in the intergenic region between VCA0543-VCA0544. | Soler-Bistué et al. 2015 |
| Parental MuGENT | N16961::mTn*7hapR^+^* Δ*lacZ*. Er^S^, Gn^R^ and Cm^S^. The Spec^R^ cassette was inserted in a neutral region between VC1902-VC1903. | This work |
| *flrA** | Parental MuGENT *flrA*^FS^ Δ(C)_5_ at base number 2,163,111. Spec^R^ cassette inserted in neutral region between VC1902-VC1903. | This work |
| *flrB** | Parental MuGENT *flrB** ΔGCGAGTG at base number 2,162,019 . Spec^R^ cassette inserted in neutral region between VC1902-VC1903. | This work |
| *mgtE** | Parental MuGENT *MgtE** point mutation, G for C at chromosome II position 765560. The Spec^R^ cassette was inserted in a neutral region between VC1902-VC1903. | This work |
