## Supplementary material for "Chromosomal position of ribosomal protein genes impacts long term evolution of *Vibrio cholerae*": Table S2

**Table S2.** Full list of primers used in this study:

| Primer name | Sequence (5’-3’) | Target |
| --- | --- | --- |
| rctB_qPCR_F | GAAGTCTCTGAAGCCGCGATTG | *rctB* |
| rctB_qPCR5-R | CCTCTTCGGAAGCGGTGATG | *rctB* |
| Tgt4-1 | ATCGGTTGCGTCACCAAATG | Superintegron (VCA0543) |
| Tgt4-4 | GCATTCGATTCGTCGTTTGG | Superintegron (VCA0544) |
| MUGENTT_FlrAFS_Seq1 | TTACGAGGAGAGGCATTGTG | VC2137/*/flrA* |
| MUGENTT_ FlrAFS_1 | TCGGTCTTCGCCACTTTATC | VC2137/*/flrA* |
| MUGENTT_ FlrAFS_2 | CAAGCGTTAGAAGCGCAATAGTGGCGAGAGCGGCAGAC | VC2137/*/flrA* |
| MUGENTT_ FlrAFS _3 | TGCCGCTCTCGCCACTATTGCGCTTCTAACGCTTGGT | VC2137/*/flrA* |
| MUGENTT_ FlrAFS _4 | TTGACTGCCGCTGAAGTGGG | VC2137/*/flrA* |
| MUGENTT_ FlrA FS_Seq2 | CGCAAATGCTCGATGATGTG | VC2137/*/flrA* |
| F_Verif_FlrAdelC | TCTCGCCACTATTGCGCTTC | VC2137/*/flrA* |
| R_Verif_ FlrAdelC | CCATATCCCGTTGCTGGTTG | VC2137*/flrA* |
| MUGENTT_FlrB_Seq1 | TATCCGGCTCACGTACCCAG | VC2136/*flrB* |
| MUGENTT_FlrB_1 | CATCGGCCACAGTCTATTACC | VC2136/*flrB* |
| MUGENTT_FlrB_2 | CAAAATCATGGAACCCTTTAAGGTACAGGCCTTGGCTTA | VC2136/*flrB* |
| MUGENTT_FlrB_3 | TAAGCCAAGGCCTGTACCTTAAAGGGTTCCATGATTTTGT | VC2136/*flrB* |
| MUGENTT_FlrB_4 | CCTTGTTGGGCAAAGCATGG | VC2136/*flrB* |
| MUGENTT_FlrB_Seq2 | CCATATCCCGTTGCTGGTTG | VC2136/*flrB* |
| R_Verif_FlrB | ATCTCGCTGCGCAATGGACG | VC2136/*flrB* |
| F_WT_FlrB | AGGCCTGTACCTTGTGAGCGAGTG | VC2136/*flrB* |
| MUGENTT_MgtE_Seq1 | CACACCTGACACCCTAGATG | VCA0818/*mgtE* |
| MUGENT_MgtE_1 | TTTCTGTACGCCCTTGATGC | VCA0818/*mgtE* |
| MUGENT_MgtE_2 | AAGCCGACCACATCGGTCACGGGTGGTGAG | VCA0818/*mgtE* |
| MUGENT_MgtE_3 | CAGGTTCGGTTATTCTCACCACCCGTGACC | VCA0818/*mgtE* |
| MUGENT_MgtE_4 | GCAGCGTTGTAGCCGTAATG | VCA0818/*mgtE* |
| MUGENT_MgtE_Seq2 | GCCGCAGACTCTTTGTCTAC | VCA0818/*mgtE* |
| F_Verif_MgtE | TTCGGTTATTCTCACCACTC | VCA0818/*mgtE* |
| R_Verif_MgtE | ACTCGGAGAGGATCAGATAC | VCA0818/*mgtE* |
