## Supplementary material for "Chromosomal position of ribosomal protein genes impacts long term evolution of *Vibrio cholerae*": Figure S1

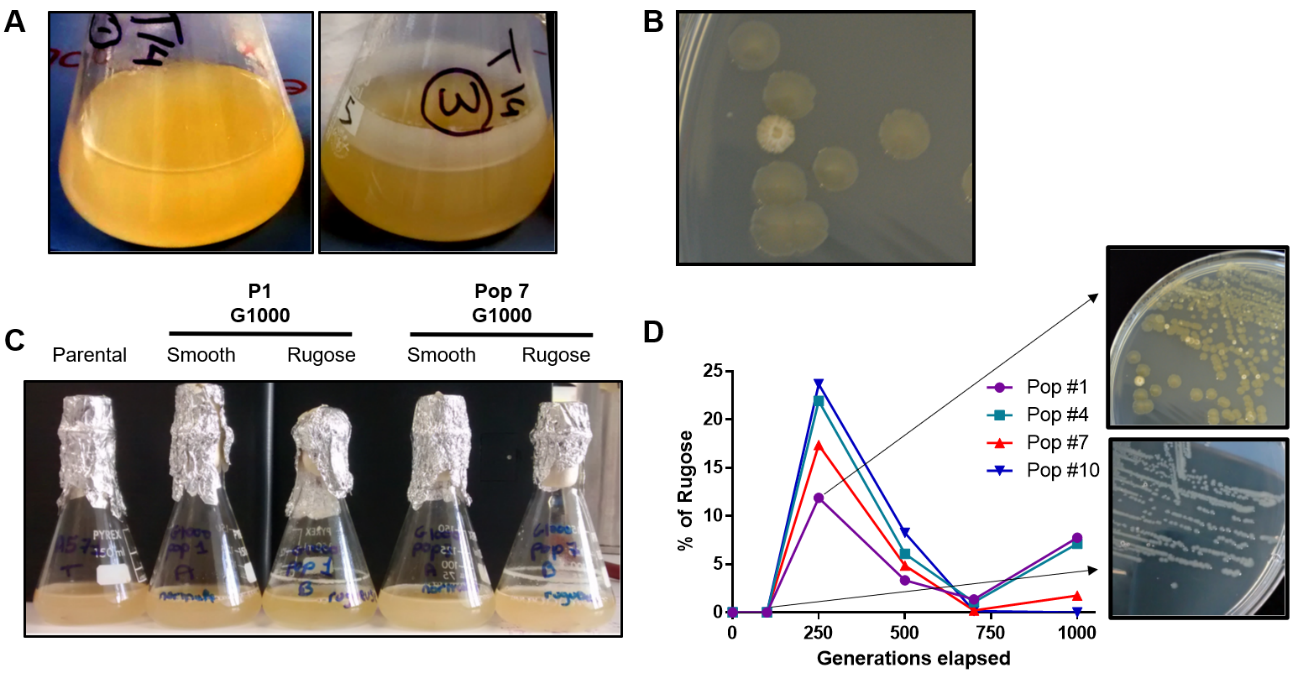

**Figure S1: Qualitative characterization of biofilm formation during the LTEE.** **A)** General aspect of a population displaying strong biofilms (P3) and a population lacking this trait (P1) forming after 114 generations. **B)** Macroscopic aspect of a smooth and a rugose colony in LB agar. **C)** Erlenmeyer cultures of the Parental strain at G0, a rugose and a smooth clone from P1 and P7 isolated at G1000. R strains showed strong biofilms in the liquid air-interphase absent in the other clones. **D)** The proportion of rugose clones along the generations is shown for 4 representative populations. The inset shows a representative photograph of P1 straked on LB agar with at maximum (upper panel, G250) and a minimum (lower panel, G100) proportion of R clones.
