## Supplementary material for "Chromosomal position of ribosomal protein genes impacts long term evolution of *Vibrio cholerae*": Figure S2

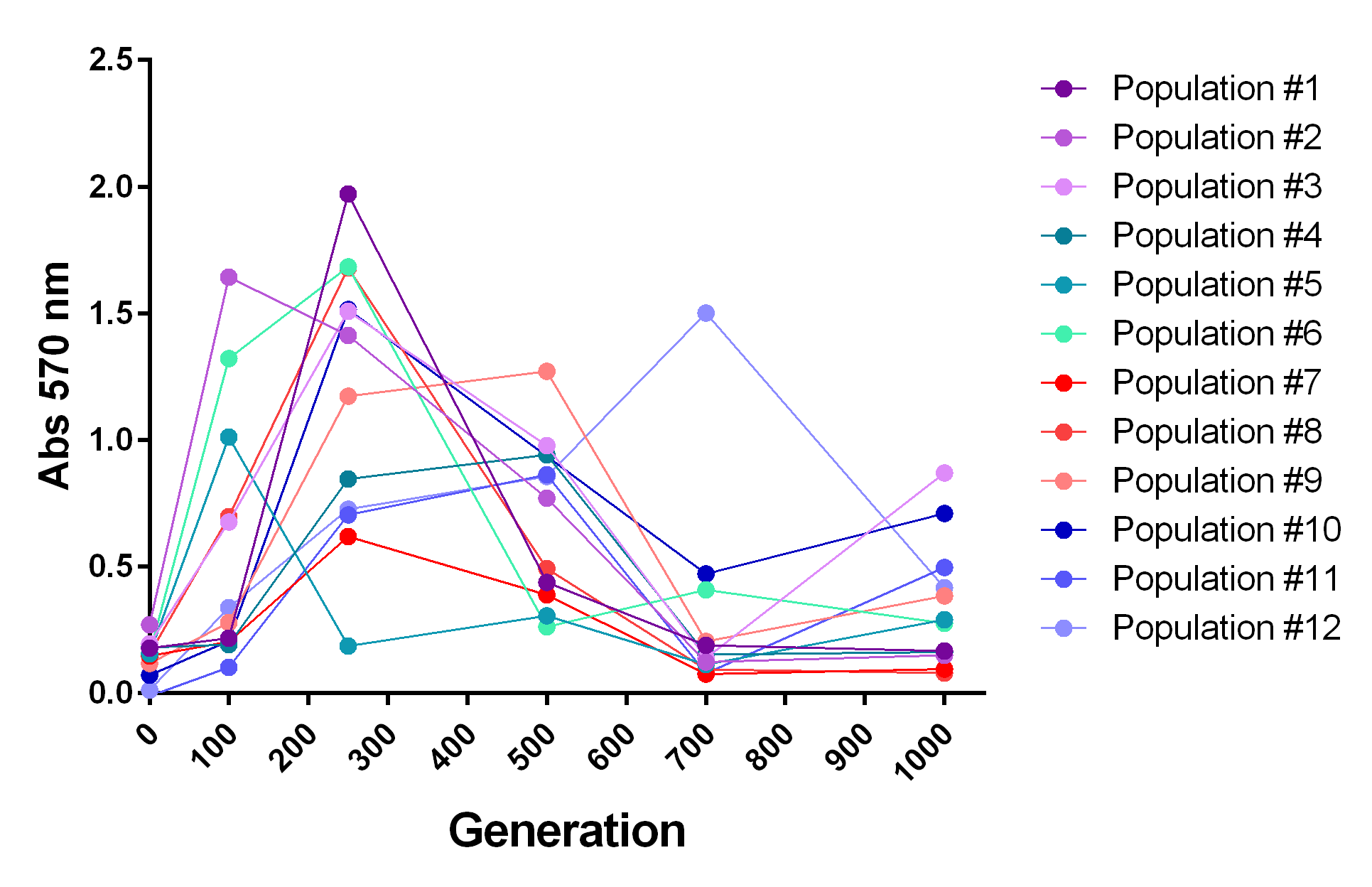

**Figure S2: Quantitative characterization of biofilm formation along the LTEE.** Crystal violet staining assays for all populations along the LTEE. The parental (Violet nuances, P1-P3) and S10Tnp-35 (Cyan nuances, P4-P6), S10Tnp-1120 (Red nuances, P7-P9) and S10TnpC2+479 (Blue nuances, P10-P12). Data averaged the means from three independent experiments.
