## Supplementary material for "Chromosomal position of ribosomal protein genes impacts long term evolution of *Vibrio cholerae*": Figure S3

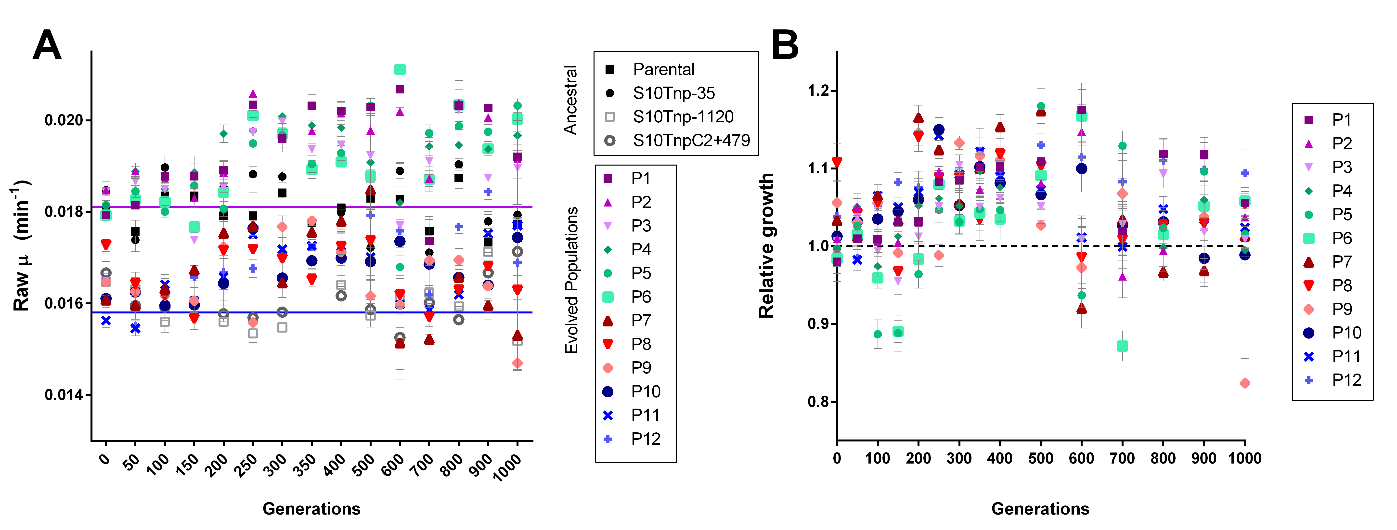

**Figure S3:** **Growth rate evolution along the generations elapsed. A)** At every time point, the populations from the frozen fossil record and each of the ancestral strains were subjected to automated growth curve. The graph shows the average at least two experiments done by triplicate of µ ± SD as a function of the generations elapsed. The average µ from the Parental (Violet) and (Blue) is shown as dotted lines for reference. The trend is similar to Figure 3 showing that relativizing growth parameters to the parental strain did not alter the result of the experiment. **B)** The obtained µ for each population was divided by the growth rate of its own ancestral strain. Then relative growth was plotted as a function of generations. The dotted line indicates 1 which represents no change with respect to the ancestral strain founding the population.
