## Supplementary material for "Chromosomal position of ribosomal protein genes impacts long term evolution of *Vibrio cholerae*": Figure S4

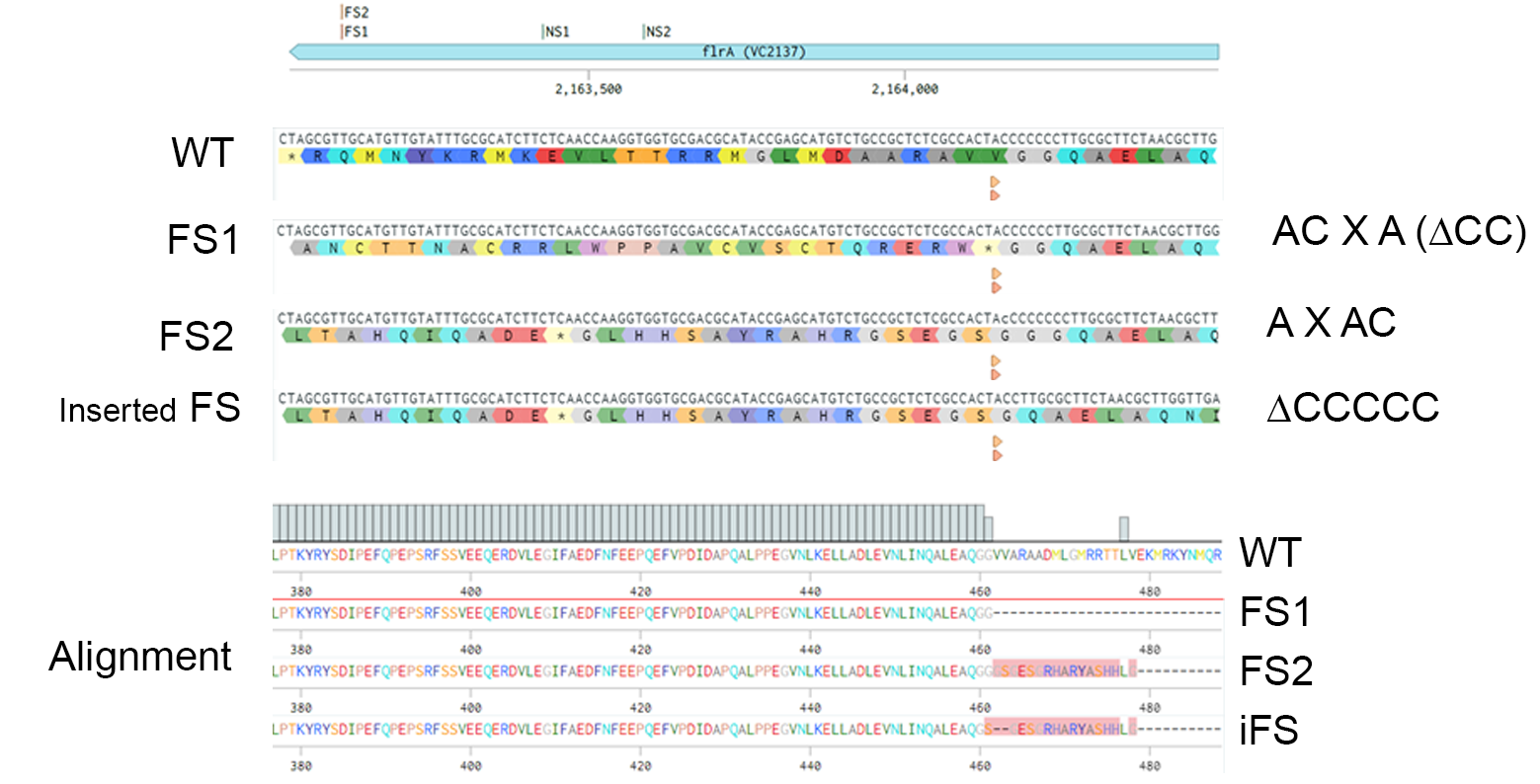

**Figure S4: Mutations characterized and generated in *flrA*.**  **Upper panel:** *flrA* gene map in scale indicating the frameshift (FS) and non-synonymous mutations (NS1) found along the EE. **Middle panel:** DNA the wild type (WT) gene, FS1, FS2 and the generated FS (inserted FS) and the resulting translation. **Lower panel:** alignment of the translated sequences showing similar early protein termination.
